# A control theoretic model of adaptive behavior in dynamic environments

**DOI:** 10.1101/204271

**Authors:** Harrison Ritz, Matthew R. Nassar, Michael J. Frank, Amitai Shenhav

## Abstract

To behave adaptively in environments that are noisy and non-stationary, humans and other animals must monitor feedback from their environment and adjust their predictions and actions accordingly. An under-studied approach for modeling these adaptive processes comes from the engineering field of control theory, which provides general principles for regulating dynamical systems, often without requiring a generative model. The proportional-integral-derivative (PID) controller is one of the most popular models of industrial process control. The proportional term is analogous to the “delta rule” in psychology, adjusting estimates in proportion to each successive error in prediction. The integral and derivative terms augment this update to simultaneously improve accuracy and stability. Here, we tested whether the PID algorithm can describe how people sequentially adjust their predictions in response to new information. Across three experiments, we found that the PID controller was an effective model of participants’ decisions in noisy, changing environments. In Experiment 1, we re-analyzed a change-point detection experiment, and showed that participants’ behavior incorporated elements of PID updating. In Experiments 2-3 we developed a task with gradual transitions that we optimized to detect PID-like adjustments. In both experiments, the PID model offered better descriptions of behavioral adjustments than both the classical delta-rule model and its more sophisticated variant, the Kalman filter. We further examined how participants weighted different PID terms in response to salient environmental events, finding that these control terms were modulated by reward, surprise, and outcome entropy. These experiments provide preliminary evidence that adaptive behavior in dynamic environments resembles PID control.

---

To behave adaptively, we must adjust our behavior in response to the dynamics of our environment (Ashby, 1956; Pezzulo & Cisek, 2016). Achieving this goal requires us to collect feedback about the outcomes of our recent actions, and incorporating this feedback into decisions about how to adjust future actions. Within research on learning and decision-making, a popular approach for achieving this feedback-based control is the *delta-rule model*^1^ (Δx = *αδ*; Widrow & Hoff, 1960; cf. Maxwell, 1868). This model adjusts expectations (x) proportionally to the discrepancy between observed and predicted outcomes (i.e., *prediction error*, *δ*), depending on the *learning rate* (*α*). While there is substantial cross-species evidence for delta-rule controlled behavior (e.g., Rescorla & Wagner, 1972; Mirenowicz & Schultz, 1994; Garrison, Erdeniz, & Done, 2013), this algorithm has major limitations. The delta-rule is sensitive to any noise that will cause persistent errors, either leading to oscillatory behavior (at a high learning rate) or a sluggish response (at a low learning rate; Rumelhart, Hinton, & Williams, 1986; Aström & Murray, 2010). However, one of the greatest limitations of this algorithm is that it performs poorly in environments that are non-stationary (i.e., that change discontinuously over time; Pearce & Hall, 1980; Aström & Murray, 2010).

More elaborate feedback control mechanisms have been developed for use in various industrial applications within a branch of engineering that studies the regulation of dynamical systems called *Control Theory*. Many control theoretic algorithms augment the basic delta rule with additional control terms that greatly improve accuracy, stability, and responsivity. The most popular variant of these control theoretic models is the popular proportional-integral-derivative (PID) controller (Fig. 1). This model is simple, accurate, and robust, with response properties that have been well-characterized over the last century (Franklin, Powell, & Emami-Naeini, 1994; Aström & Murray, 2010). The PID controller takes the error from a reference signal as input, and it outputs a control signal consisting of a linear combination of control signals proportional to the error (P-Term), the integral of the error (I-Term), and the derivative of the error (D-Term; Fig. 1). These three terms minimize deviations from the reference based on errors in the present, past, and expected future, respectively.

**Figure 1.**
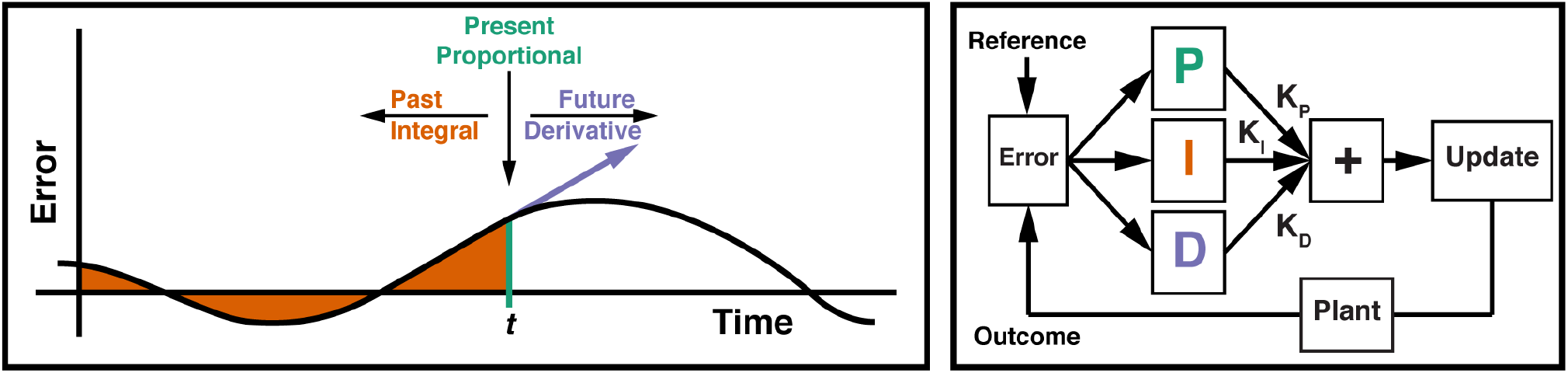
**Left:** The Proportional-Integral-Derivative (PID) Controller uses the current error (P-Term), the integral of the error (I-term), and the derivative of the error (D-Term), in order to provide compensation based on the present, past, and future, respectively. **Right:** The PID controller specifies a control signal based on the weighted sum of the PID terms, with each term weighted by their respective gain. Similar to a thermostat-controlled furnace, the plant implements the control signal, moving the measured processes closer to the reference (e.g., the desired temperature). Left panel adapted from (Aström & Murray, 2010), permission pending.

Proportional control (cf. delta-rule control) directly minimizes deviation from the reference and is often the primary driver of the control process. Integral control provides low-frequency compensation for residual steady-state errors, allowing the controller to reduce noise and track gradual changes in the environment. Derivative control provides high-frequency compensation that increases stability, dampening control adjustments when the controller is approaching the reference and increasing adjustments when the reference or environment changes suddenly (see: Aström & Murray, 2010). Intuitively, integral control provides low-frequency compensation by combining several time-points, whereas derivative control providence high-frequency compensation by tracking the instantaneous change. Here we test whether this popular model of industrial control can account for adjustments in human behavior within a dynamic environment.

PID control has algorithmic properties that make it useful for most control systems. For instance, relative to algorithms that require an explicit representation of task dynamics, PID can provide an effective, and computationally cheaper, model-free alternative to adjusting cognitive or behavioral processes over time, particularly for natural environments that require particularly complex world models. Moreover, convergent evidence suggests that the PID algorithm may help account for the variety of feedback-related findings observed in humans and other primates. Behavioral and neural correlates of feedback-controlled choice provide preliminary evidence that participants transform decision-relevant variables in a manner predicted by the PID algorithm. Consistent with proportional control, there is substantial evidence that participants adjust their behaviors based on recent errors or conflict (Rescorla & Wagner, 1972; Lau & Glimcher, 2005; Rabbitt, 1966; Gratton, Coles, & Donchin, 1992; Ullsperger, Danielmeier, & Jocham, 2014), with corresponding signals observed most prominently in the striatum and anterior cingulate cortex (ACC; Mirenowicz & Schultz, 1994; Garrison, Erdeniz, & Done, 2013; Niki & Watanabe, 1979; Ito, Stuphorn, Brown, & Schall, 2003; Matsumoto, Matsumoto, Abe, & Tanaka, 2007; Kennerley, Walton, Behrens, Buckley, & Rushworth, 2006; Seo & Lee, 2007; Smith et al., 2015). Previous work has found that people are also sensitive to the extended history of errors or conflict (Laming, 1968; Logan & Zbrodoff, 1979; Botvinick, Braver, Barch, Carter, & Cohen, 2001; Aben, Verguts, & Van den Bussche, 2017; Alexander & Brown, 2015; Bugg & Crump, 2012; Wittmann et al., 2016), with proposals that this specifically involves integrating over recent errors (Alexander & Brown, 2015; Wittmann et al., 2016). Accordingly, experiments have found neural signals in the prefrontal cortex and ACC that reflect this feedback history (Carter et al., 2000; Blais & Bunge, 2010; Bernacchia, Seo, Lee, & Wang, 2011; Wittmann et al., 2016), and recent models of the ACC have emphasized the role that integrative, recurrent activity in this region play in executive control (Wang, 2008; Hunt & Hayden, 2017; Shahnazian & Holroyd, 2017). Finally, consistent with derivative control, prior work has found that participants track the environmental rate of change when making decisions, with associated neural correlates in the anterior prefrontal cortices and ACC (Behrens, Woolrich, Walton, & Rushworth, 2007; Jiang, Beck, Heller, & Egner, 2015; Bernacchia et al., 2011; Kovach et al., 2012; McGuire, Nassar, Gold, & Kable, 2014; Wittmann et al., 2016). While some of these results have been attributed to participants’ representations of environmental dynamics (e.g., Behrens et al., 2007; McGuire et al., 2014; Jiang et al., 2015), PID control may offer a parsimonious account of these behaviors.

Despite the success of the PID model as a simple and effective algorithm for implementing control in other fields, and suggestive evidence for relevant neural signatures in circuits involved in adaptive control, PID has yet to be formally tested as a model of human adaptive behavior. In the current set of experiments, we directly tested whether a PID model can describe human performance in adaptive learning tasks. In Experiment 1, we re-analyzed a recent study that examined predictive inference in an environment with discrete change points. Behavior on this task confirmed key predictions of the PID model, but was limited in its ability to adjudicate between candidate models. Informed by our findings in Experiment 1, for Experiments 2-3 we developed a novel task that was optimized for PID control, using gradual rather than sudden change-points. We found that the PID model was a strong predictor of participants’ choices in both experiments. Experiment 3 replicated the predictive power of our model, and further examined whether participants dynamically adjust their control terms based on rewards, surprise, and outcome entropy. Across these tasks, participant performance was consistent with key predictions of the PID model, demonstrating that this simple model provides a promising account of adaptive behavior.

## Experiment 1

The PID model is designed to adapt the behavior of a system in response to changes in the environment. We therefore began by testing whether this model could explain behavioral adjustments in an existing change-point detection task, one that was designed to assess how humans can adapt their learning rate to uncertain and volatile outcomes (McGuire et al., 2014). In this experiment, participants predicted where a target stimulus would appear (horizontal location on the screen), and then received feedback about where the true location of the outcome had been (See Fig. 2A). Outcome locations were normally distributed around a mean, and the mean of this distribution changed suddenly throughout the experiment (change points), according to a predetermined hazard rate. This task allows us to measure participants’ choices and feedback in a continuous space with high precision, making it desirable for studying PID control. We can therefore use the PID model to predict trial-to-trial adjustments in participant behavior (predicted locations) based on their history of error feedback. In other respects, this task is not ideally suited for testing our model: the dramatic changes in target distributions may ‘reset’ adaptive control processes (Bouret & Sara, 2005; Karlsson et al., 2012; Tervo et al., 2014), and so this experiment serves as a preliminary test of our hypothesized control dynamics. We will address these concerns in Experiments 2-3.

**Figure 2.**
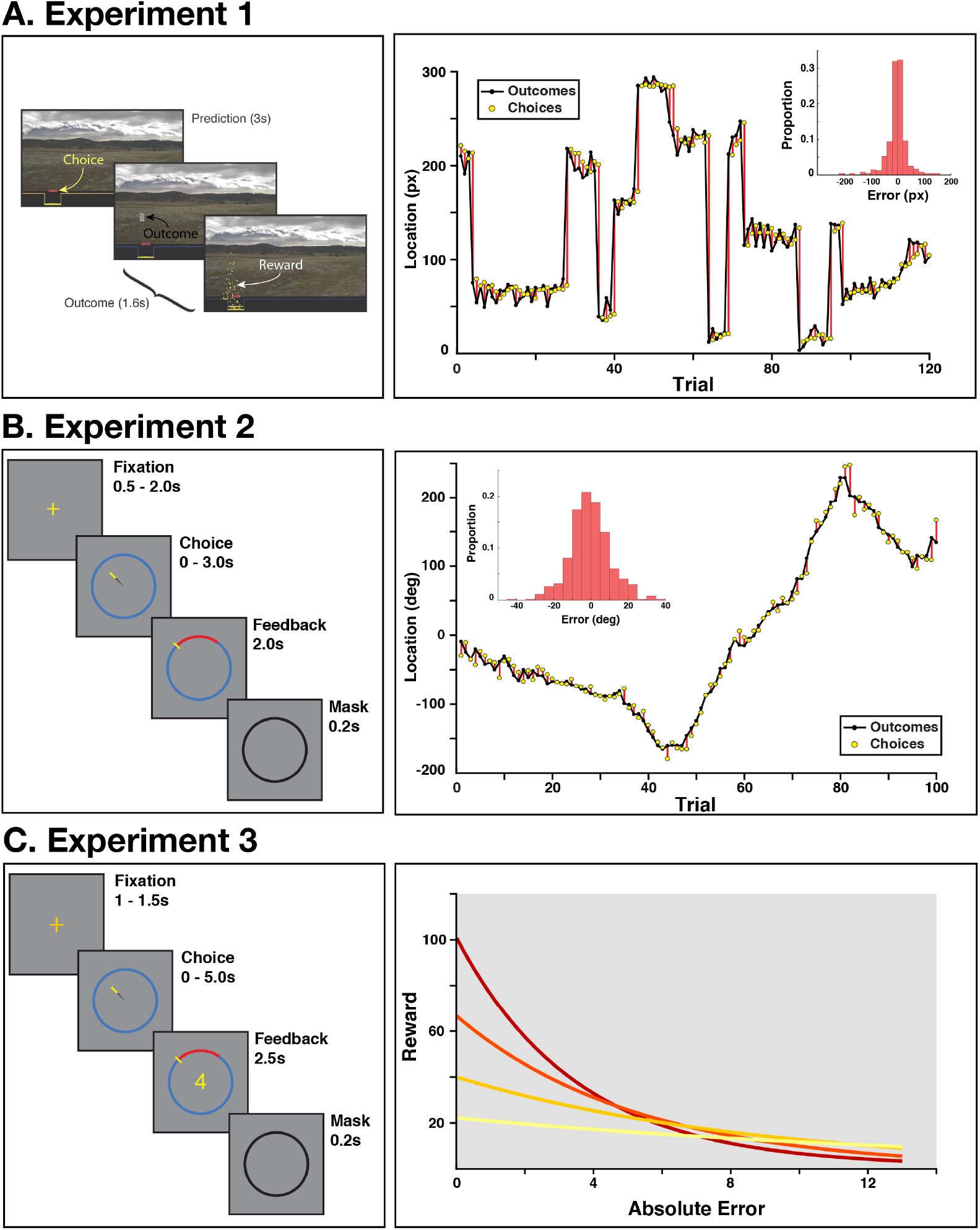
Experimental tasks. **A) Left:** On each trial of Experiment 1, participants selected a horizontal location with a joystick and were then shown the correct location. On a random subset of trials, participants received performance-contingent rewards (shown as gold coins). Figure adapted from McGuire et al., 2014, permission pending **Right:** A representative block of trials from an example participant. The mean correct location was stable for a variable number of trials, and then was uniformly resampled. Inset histogram: The distributions of errors for this participant across their session. **B) Left:** Participants in Experiment 2 selected a location on the circle with their mouse, and then were shown the correct location. **Right:** A representative block of trials, demonstrating that the mean correct location changed gradually over time. As seen in the histogram for an example participant (inset), the gradual changes in location for this task resulted in error distributions that were less peaked than in Experiment 1 (compare Panel A inset). **C) Left:** Experiment 3 was identical to Experiment 2, but participants were rewarded based on their accuracy, according to one of four reward-error functions. They were informed of the current reward mode during Fixation, and during Feedback they received the reward corresponding to their accuracy on that trial (conditional on the current reward mode). **Right:** Error-reward slopes for the four reward modes.

## Methods

### Participants and Procedure

Experiment 1 consisted of a re-analysis of a change-point task used by McGuire and colleagues (2014). Briefly, thirty-two participants (17 female, mean(SD) age = 22.4(3.0)) selected a location on a line using a joystick, and then were shown the correct location for that trial. On a random subset of trials, participants were rewarded according their accuracy, dissociating the trial value (which depended on whether the trial was rewarded) from errors. Target locations were drawn from a Gaussian distribution with a variable mean and variance. The mean was stable for three trials, and then on a weighted coin flip (hazard rate: 0.125) was uniformly redrawn from the line coordinates; the variance alternated between high and low levels across blocks. Participants performed 160 training trials followed by 4 blocks of 120 trials during fMRI. Two participants were excluded for having an inconsistent number of trials per block, leaving 30 participants for the final analysis. See McGuire et al. (2014) for additional details.

### Lagged Regression Analysis

A critical prediction of the PID model is that a participant’s updates (i.e., the change in their location guesses) should depend on their history of feedback. While a delta-rule model predicts that only the error on the current trial would directly influence updates, a PID controller integrates errors over a longer history, enabling the controller to correct for a consistent bias in errors. Integral control will manifest as an exponentially decaying influence over all previous trials, whereas derivative control will place a positive weight on the current trial, and a negative weight on the *t*-1 trial. These two terms make different predictions for the *t*-1 trial: integral control will place a high weight on this trial, whereas derivative control places a lower weight on *t*-1 than it does on earlier trials.

To measure the independent contribution of each trial’s feedback in the recent past, we used a simple lagged regression analysis to test how prediction updates (change in predicted location from the current to next trial) depended on the errors from the current and 10 previous trials (*u*_*t*_ ˜ 1 + *e*_*t*_ + *e*_*t*−1_ + … *e*_*t*−10_; Wilkinson notation). We assessed the influence of pervious trials’ feedback by testing whether the sum of previous trials’ betas was significantly different from zero, using a nonparametric sign-randomization test at the group level (comparing the observed results to a null distribution that we generated by randomly assigning positive or negative signs to each participant’s summed betas). Throughout the paper, all randomization tests used 10^5^ simulations, and all statistical tests were two-tailed with α = .05.

### PID Model

The PID algorithm controls a system in order to maintain a desired reference signal (Fig. 1). It takes as input the signed error relative to this reference (*et* = reference - output), and produces a control signal (*u_t_*) that specifies the adjustment for the next time point (here, the next trial). The control signal is defined by a linear combination of three terms: the P-term (reflecting the error), the I-term (reflecting the leaky integration of the error), and the D-term (reflecting the derivative of the error). Each of these terms was weighted by its own gain parameter (*K_P_*, *K_I_*, *K_D_*). For trial *t*, the control signal (*u_t_*) was generated by transforming the error(*e_t_*) as follows:

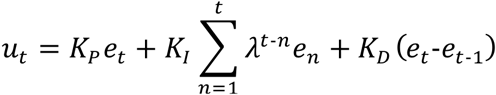

where λ represents a memory persistence parameter, with larger values leading to longer retention. On the first trial of a block, the I-Term is *e_t_*, and the D-Term is 0, producing a control action similar to proportional control. In the following tasks, *u_t_* was defined as the difference in the choice location between trial *t+1* and trial *t* (hereafter, the ‘update’), and *e_t_* was the difference between the correct location and the chosen location. While PID is not traditionally a state-estimation algorithm, it can serve this function by regulating performance to maintain a desired accuracy (i.e., no error), such as occurs in autoencoder learning systems (Denève, Alemi, & Bourdoukan, 2017).

### PID Model Fit

We used each participant’s timecourse of errors within each block to generate hypothesized P, I, and D values based on the raw errors, the integral of the errors, and the first derivative of the error, respectively. Our regression model consisted of an intercept and the three PID terms (*u ~1* + *P* + *I* + *D*), and we fit this model with iteratively reweighted robust regression (using MATLAB’s fitlm function; bisquare weighting factor), to minimize overfitting to the outliers that can occur when participants make scalar responses. Fit statistics were generated based on the non-reweighted residuals from the robust model, in order to avoid undue bias in favor of complex models.

Because the λ parameter (memory persistence) interacted with the identifiability of our PID terms when estimated jointly (e.g., when λ = 0, P and I are identical), we chose to fit this single term at the group rather than individual level. We fit λ with a grid search (range: [0.5-1], increments of 0.001), using median *R^2^* across participants as our measure of fit (normalizing individual differences in update variability). Regression models were estimated at the individual level, and regression weights were tested for deviance from zero at the group level with a sign-randomization test (see above).

We compared the P (i.e., delta-rule), PI, PD, and PID models, as these are the most common instantiations of the PID algorithm. To compare model performance, we used each participant’s Akaike’s Information Criterion (AIC; Akaike, 1983), an index of goodness-of-fit that is penalized for model complexity^2^. We compared the AIC at the group level using Bayesian model selection (Rigoux, Stephan, Friston, & Daunizeau, 2014), quantifying the Bayesian omnibus risk (BOR; probability that all models are equally good across the population) and each model’s protected exceedance probability (PXP; the probability that this model is more frequently the best fit than any of the competing models, controlling for the chance rate). BOR tests whether there is an omnibus difference between models, whereas PXP describes which models fit the best.

### PID Controller Simulations

To better understand the expected range of behavior under our candidates models, we simulated delta-rule and PID controllers performance for the outcome histories that each participant encountered during the experiment. We used a restricted maximum likelihood estimation procedure (MATLAB’s fmincon) to determine the values of the PID gains, λ, and choice bias (i.e., intercept) that perform best given the outcomes of each participant’s task. We then compared these best-performing delta-rule and PID gains to the gains estimated from participants’ behavior.

We also tested whether our fitted model would produce the same pattern of behavior that we measured with our lagged regression. We simulated an ideal observer that used each participant’s estimated PID parameters and outcome history in order to generate a sequence of updates, and then fit our lagged regression to this simulated behavior, separately for the P, PI, and PID candidate controllers. This analysis allows us to qualitatively determine the extent to which the PID model can act as a generative model of participants’ decision-making behavior (Gelman, Meng, & Stern, 1996; Nassar & Frank, 2016). If participants are using PID control, then simulated updates from a PID controller should similarly weight the feedback received over previous trials.

## Results

### Model-Agnostic Analysis

To identify the degree to which behavioral adjustment were influenced by recent feedback, we regressed participants current and previous errors on their update. We found that the current trial was the strongest predictor of the current update, but that updates were also influenced by feedback from leading trials (Fig. 3A). The sum of leading trials’ betas was reliably less than zero (Mean(SD) summed beta: −0.040(0.064), *p* = .00017). This suggests that while immediate feedback was the most influential factor for participants’ updates, they also incorporated an extended history of error feedback into their adjustments. While the current trial had a positive influence on updates, these previous trials instead had a negative influence on the current update, potentially compensating for the extreme errors that participants made at a change-points (see Fig. 2A).

**Figure 3.**
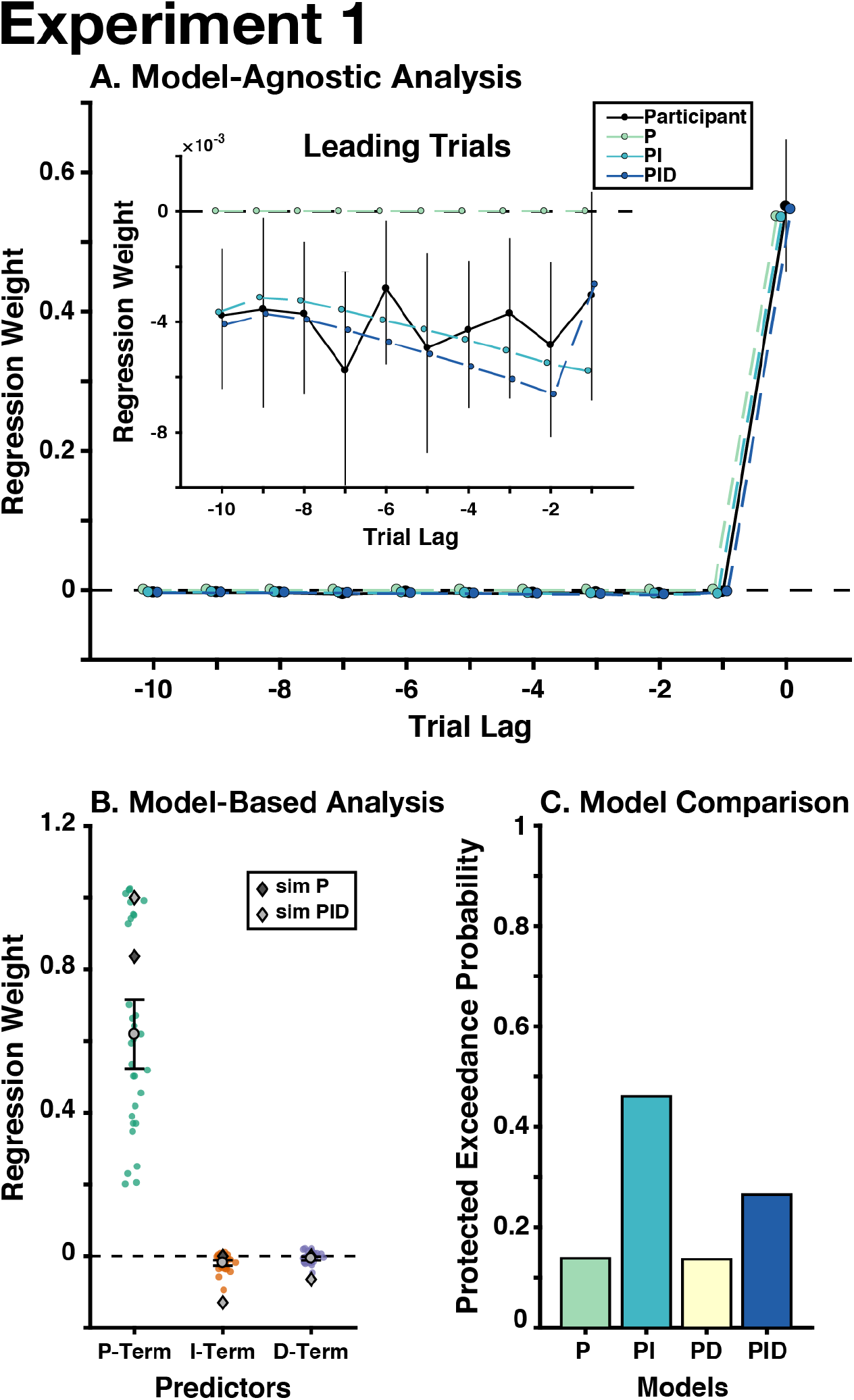
Experiment 1 Results. **A)** For our model-agnostic analyses, we regressed the errors that participants made on the current and ten leading trials on their current update (black: participants’ regression weights). Next, we used the PID parameters estimated in our regression analysis to generate behavior from P, PI, and PID controllers, and fit our lagged regression to this simulated behavior (colored lines). Inset: the regression weights from only the leading trials (i.e., before the current trial), controlling for the effect of the current trial. **B)** For our model-based analyses, we regressed the trial-wise P-, I-, and D-Terms on participants’ updates, and found that all three terms were significantly different from zero. Colored circles indicate individual participants’ regression weights. Dark gray diamonds indicate the mean gains from the best-performing delta-rule models, based on each participant’s outcome history; light gray diamonds indicate the mean gains from the best-performing PID controllers. **C)** We used Bayesian model selection to adjudicate between our candidate models, finding that the PI model best explained the data, albeit with moderate support (see text). Error bars throughout indicate mean and between-participant bootstrapped 95% confidence intervals.

To verify that our model can generate performance that captures the behavior observed in this task, we simulated behavior on this task using parameters estimated for our PID model and reduced versions thereof (P, PI). We then performed the same lagged regression on these simulated data that we used on real data (Fig. 3A). As expected, we found that the simulated PI and PID models captured the influence of leading errors, unlike the P-only model (which predicts that there should be no influence of leading errors).

### PID Model Fit

We first performed a search to identify the PID gains that optimized task performance (minimizing mean squared prediction error) for the outcome sequence observed by each participant. We found that the optimal PID gains were all reliably different from zero (Mean(SD) PID Gain: *K_P_* = 1.0(0.18), *K_I_* = −0.16(0.12), *K_D_* = −0.070(0.077), λ= 0.85(0.071). Consistent with the lagged-regression analysis, the optimized integral and derivative gains were negative.

Fitting our PID model to participants’ updates, we found that the best-fit models accounted for a substantial amount of this variance (median *R*^2^ = 0.92), with parameters for all terms being significantly different from zero (Mean(SD) standardized betas: *β_P_* = 0.62(0.27), *p* ≤ 10^−^5; *β_I_* = −0.019(0.022), *p* ≤ 10^−^5; *β_D_* = −0.0068(0.016), *p* = 0.022; see Fig. 3B). The group-level λ (memory persistence) was also quite high (0.9430), suggesting that participants retained a great deal of information regarding past feedback. Participants’ estimated gains qualitatively resembled the gains produced by the simulated PID controller, sharing the same sign and approximate magnitude (compare gray diamonds and circles in Fig. 3B).

We used Bayesian model selection to compare the fit of each model (PXP), and test whether there was an omnibus difference between models (BOR). Of these three models, we found that the PI model had the highest protected exceedance probability (PXP_P_ = 0.14, PXP_PI_ =0.46, PXP_PD_ = 0.14, PXP_PID_ = 0.26; Fig. 3C), but that there is altogether insufficient evidence to support one model over another (BOR = 0.55, providing roughly equal evidence that the models are the same or different). These data therefore do not allow us to rule out the possibility that a simple delta-rule (P-only) model parsimoniously accounts for participant behavior. The PID models did not predict behavior better than the Bayesian change-point model in the original publication (original median *R*^2^ = 0.97; McGuire et al., 2014), which incorporated information about the generative structure of the statistical environment.

## Discussion

We found preliminary evidence that participants performing a change-point detection task are influenced by their history of error feedback, consistent with the predictions of a PID controller. Participants’ updates could also be predicted from the integral and derivative of their errors. Despite these promising indications of PID control, we were unable to confidently differentiate between candidate models. Furthermore, this model did not explain behavior better than the change-point detection model from this original experiment.

While this experiment offers mixed evidence in favor of the PID algorithm, this may be because this task was designed for change-point models, with sudden, dramatic shifts in the outcome distribution. These change-points introduces extreme errors that participants might treat categorically differently from normal prediction errors, evoking a ‘reset’ in their decision process or causing the representation of a different context (Bouret & Sara, 2005; Courville, Daw, & Touretzky, 2006; Nassar, Wilson, Heasly, & Gold, 2010; McGuire et al., 2014; O’Reilly et al., 2013). This experiment also involved a training paradigm designed to make participants aware of the generative structure of the task, an advantage not typically afforded in the real world or exploited by PID control systems. With these concerns in mind, we developed a novel adaptive learning task in which outcomes changed smoothly over time, encouraging participants to treat outcomes as arising from a single, changing context. Participants were not instructed on this generative structure explicitly, reducing the potential for the use of structured inference strategies that best characterized learning in Experiment 1.

## Experiment 2

While Experiment 1 provided promising evidence that participants use their history of feedback in a way that resembled PID control, it did not provide definitive evidence as to whether this is the best explanation for the data. However, participants may have strategically reset their predictions at extreme change-points, making it more difficult to measure history-dependent predictions. To address this, we developed a task with gradual transitions in which participants tracked an outcomes distribution whose mean linearly changed from one location to another, and whose variance changed randomly throughout the block. To make these location transitions seem more continuous, outcomes appeared along a circle rather than a straight line, thus also avoiding edge effects that can occur at either end of a screen. This design allowed us to precisely measure participants’ predictions, errors, and adjustments within an environment whose dynamics are more fluid and predictable than Experiment 1. This task was explicitly designed to emulate an environment for which a PID controller is well-suited, and specifically to maximize our power to detect differences between PID control and proportional (delta-rule) control.

## Methods

### Participants & Procedure

Twenty-nine Brown University undergraduate students (25 female; mean(SD) age = 18.6(0.83)) performed a supervised learning task in which they predicted an outcome location on a circular display (see Fig. 2B).

Participants completed 5 blocks of 100 trials in which they used a mouse cursor to guess a location on the circumference of the circle. They were then shown the correct location, with an arc indicating the magnitude and direction of their error. Participants completed 50 training trials before the main experiment. Participants had up to 3 seconds to make their guess, or else their final cursor angle would be chosen as the guess for that trial, and feedback was presented for 2 seconds. Our final analysis excluded any trials where participants did not move their cursor to the edge of the circle, as well as a subset of trials following aberrant feedback due to a technical issue (1.8% of total trials).

The target location for each trial was drawn from a Gaussian distribution over arc degrees, with a mean and standard deviation that systematically changed over time. On a weighted coin flip (hazard rate: 0.80), the distribution’s mean shifted based on a random draw from *U*(−180, 180) degrees. After the new mean was drawn, the mean would transition from the old mean to the new mean over *U*(8, 20) trials, with the means during transition trials linearly interpolated between the old and new means. The standard deviation varied independently of the mean, and was redrawn from *U*(1, 8) degrees on a weighed coin flip (hazard rate: 0.40). These task parameters were selected through simulation to maximally differentiate the performance of PID and delta-rule models. Unless otherwise indicated, methods of analysis and model selection for this study are identical to Experiment 1.

## Results

### Model-Agnostic Analysis

Regressing the current and ten leading errors onto the current update (see Experiment 1 Methods), we again found that the sum of leading errors was significantly different from zero (Mean(SD) summed leading betas: 0.27(0.31), *p* ≤ 10^−^5; Fig. 4A). This replicates the observation in Experiment 1 that participants incorporate the extended history of errors into their prediction process.

**Figure 4.**
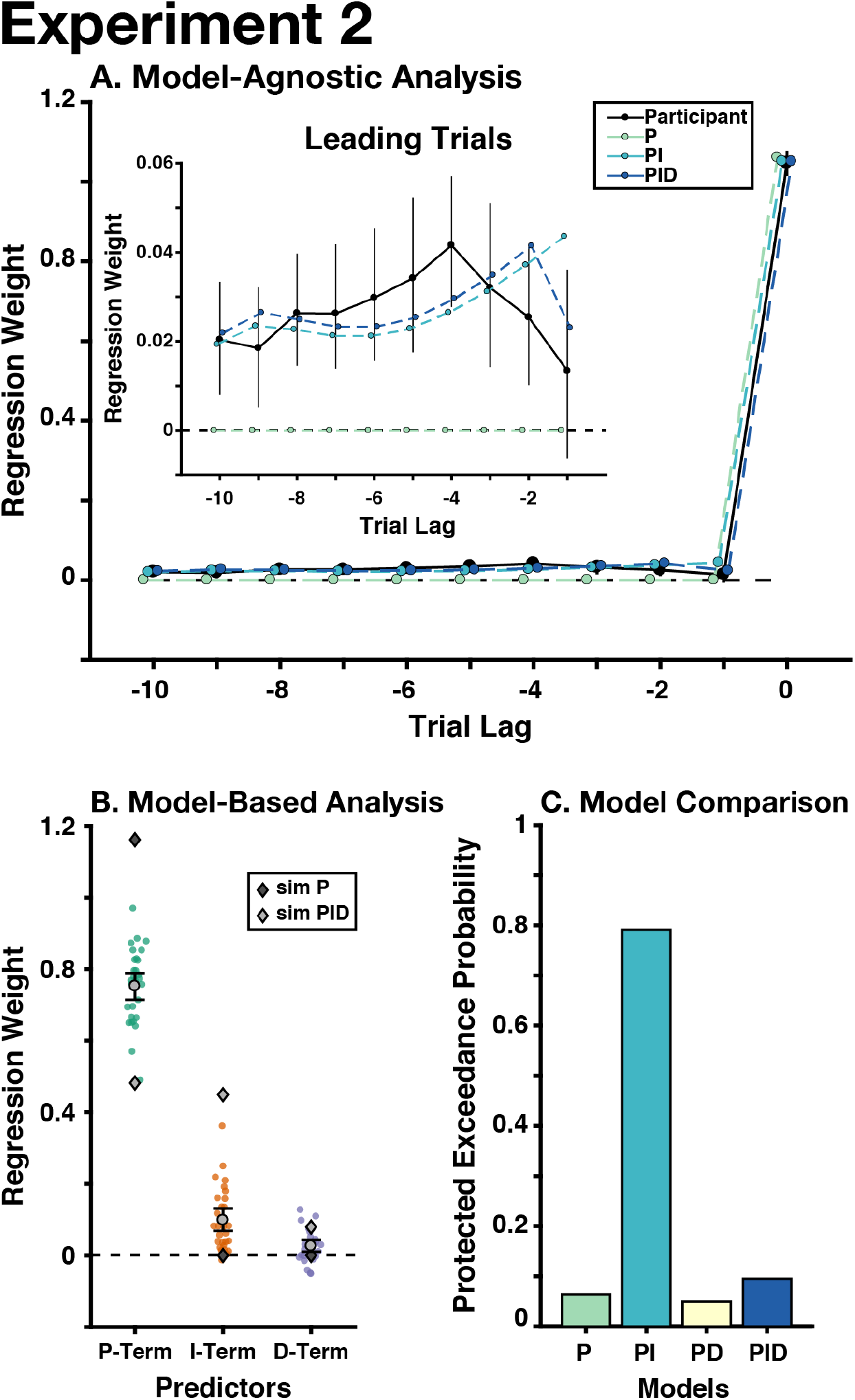
Experiment 2 Result. **A**) Participants adjusted their choices based on previous trials, unlike the predictions of a proportional controller (i.e., delta-rule model). **B**) P-, I-, and D-Terms significantly predicted participants’ updates. **C**) The PI controller best explained participants’ behavior. See Figure 3 for detailed graph legends. Error bars indicate mean and between-participant bootstrapped 95% confidence intervals.

Fitting our lagged model to behavior generated from our models produced a similar pattern of predictions as in Experiment 1: the P-only model categorically failed to capture the influence of leading errors. The PI and PID models were similar in their ability to recreate participants’ use of previous errors, however the PID model seemed to better capture participants’ weighting of recent leading errors (i.e., over the previous three trials). We examined whether this trend was reliable across participants by fitting linear and quadratic trends over trials to each participants’ leading betas. We found a significant quadratic trend (Mean(SD) quadratic trend standardized beta: −0.0054(0.009), *p* = .004), but not a linear trend (Mean(SD) linear trend standardized beta: 0.002(0.020), *p* = .39). While this finding is broadly compatible with derivative control’s non-linear weighting of previous errors, the observed trend extended further backwards in time than the previous-trial effects predicted by a simple derivative.

### PID Model Fit

The best performing PID gains were significantly different from zero (Mean(SD) gain: *K_P_* =0.48(0.13), *K_I_* = 0.45(0.085), *K_D_* = 0.080(0.066), λ = .84(0.033); all *p*s ≤ 10^−^5; Fig. 4B). Our PID model accounted for most of the variance in participants’ updates (median *R*^2^ = 0.84), and the parameters for the P, I, and D-Terms were all significantly different from zero (Mean(SD) standardized beta: *β_P_* = 0.75(0.10), *p* ≤ 10^−^5; *β_I_* = 0.099(0.088), *p* ≤ 10^−^5; *β_D_* = 0.026(0.045), *p* = 0.0034; sign-randomization test; Fig. 4B). The group-level λ was .87. Participants’ parameters were similar to the ideal PID controller, although they had a greater reliance on proportional control and a lesser reliance on integral control than was optimal.

Bayesian model selection favored the PI model (PXP_P_ = .064, PXP_PI_ = .79, PXP_PD_ = .049, PXP_PID_ = .095; Fig. 4C), although there was a moderate likelihood that models did not differ in their fit (Bayesian omnibus risk = 0.20). This was mostly due to the similarity in likelihood between the PI and PID model (excluding PID: Bayesian omnibus risk < .001, PXP_PI_ > .99). Therefore, our model selection supports the interpretation that PI control explains behavior better than the delta-rule model.

## Discussion

Using a novel variant of a change-point task, we provide strong evidence that the PI control model can usefully describe participants’ predictions. Our model-free analysis showed that participants incorporated previous errors in their adjustments, in a way that is not predicted by a proportional control model, but is well approximated by our PI model. We also found that all of the PID terms were significant predictors of participants’ updates, and that they were qualitatively similar to the gains of an ideal PID controller. Building on Experiment 1, this experiment provided strong evidence that PI was the best-fitting model. These data further support a role for control processes that extend beyond immediate errors.

These first two experiment have provided promising evidence that the PID framework predicts adaptive behaviors better than the classical delta-rule model. A striking feature of these experiments is that participants had very different estimated control gains across the two experiments, consistent with the differential gains of the best-performing PID agents. These differences suggest that participants may set their control gains in a context-specific manner, although at an unknown timescale. Popular delta-rule models have suggested that participants may in fact rapidly adjust their control gain in response to changes to the local context (Pearce & Hall, 1980). This prompted us to develop a third experiment, in order to replicate our results from Experiment 2, and test whether participants can adaptively adapt PID gains to their local contexts.

## Experiment 3

In Experiment 3, we sought to replicate the findings from Experiment 2, while at the same time manipulating the incentives for performing accurately on the task. We additionally sought to examine three factors that might influence the weights that individuals might place on each of the PID terms over the course of an experiment.

First, we examined the influence of surprise (absolute error) on these control weights, given classic findings that such surprise signals modulate learning, indicating the degree to which the environment has been learned (Pearce & Hall, 1980; see also: Hayden, Heilbronner, Pearson, & Platt, 2011; Nassar et al., 2010; McGuire et al., 2014; O’Reilly et al., 2013). Second, given evidence that learning can be influenced by uncertainty over recent feedback (Yu & Dayan, 2005; Courville et al., 2006; Nassar et al., 2010), or related estimates of volatility (Behrens et al., 2007), we examined how PID gains were influenced by an index of the outcome entropy over the past several trials. This measure of uncertainty indexes both expected uncertainty (the variance in the generative distribution) and unexpected uncertainty (changes in the mean of the generative distribution, i.e., drift), the latter of which is more dominant in our tasks.

We also examined the influence of reward on PID gains, given previous evidence that these can impact learning in a dissociable fashion from surprise or uncertainty alone (McGuire et al., 2014), and more generally that rewards may compensate for the costs of effortful control policies (Hayden, Pearson, & Platt, 2009; Padmala & Pessoa, 2011; Manohar et al., 2015; Kool, Gershman, & Cushman, 2017), including learning in particular (Hayden et al., 2009; Shenhav, Botvinick, & Cohen, 2013). For example, this could occur if integrating feedback utilizes domain-general working memory processes (Collins & Frank, 2012; 2018). Importantly, Experiments 1 and 3 were designed to de-confound reward from errors, providing us the ability to measure their influences on PID gains separately from one another and from our measure of uncertainty. In Experiment 1, performance-dependent rewards were given on a random subset of interleaved trials, whereas in Experiment 3, rewards were a nonlinear function of error that changed over time. These measures allowed us to distinguish the independent effects of surprise (absolute error) and reward on learning. For example, participants may have been motivated to perform accurately, and insofar as this motivation is further enhanced by reward, our analysis should be able to dissociate this motivation from other outcomes of error (e.g., surprise).

Finally, we compared our PID model against a popular model of adaptive learning, the Kalman filter (Kalman, 1960; Kakade & Dayan, 2002; Kording, Tenenbaum, & Shadmehr, 2007). This model performs state estimation using a delta-rule algorithm with an uncertainty-weighted learning rate. Previous experiments have found that it is a good model of behavior, and it is based on the same principles that motivated the heuristic terms in our adaptive gain analysis.

## Methods

### Participants and Procedure

Forty-seven Brown University subject pool participants (32 female; mean(SD) age = 21.3(4.07)) performed a rewarded supervised learning task (without monetary compensation). Apart from the reward manipulation, the structure of this task was similar to Experiment 2. On each trial, the reward magnitude depended on the accuracy of the participant’s guess (i.e., the absolute error between guess and outcome location; Fig. 2C). These rewards decreased exponentially with increasing error magnitude. To de-correlate rewards and errors, and to vary overall motivation to perform the task, we adjusted the steepness (mean) of this exponential (gamma) function over trials, resampling one of four possible means [1, 1.5, 2.5, 4.5] at random time points, chosen with a flat hazard rate of .20 across all trials (Fig. 2C, right). We instructed participants that these different levels of steepness defined four ‘reward modes’. The reward mode for a given trial was indicated by the color of the fixation cross (one of four colors from equally spaced locations on a heat colormap). The input (errors) to these reward functions were divided by 3.5 to approximately match the reward that these functions returned at participants’ mean performance level in Experiment 2.

Participants completed 50 training trials, followed by 6 blocks of 75 trials. On each trial, participants had up to 5 seconds to make their guess, feedback was presented for 2.5 seconds, and then the reward mode for the next trial was displayed during an ITI that was drawn from *U*(1, 1.5). At the end of each block, participants were shown the mean reward earned during that block. Our final analysis excluded any trials where participants did not move their cursor to the edge of the circle (.07% of total trials). The lagged and trial-wise regression analyses were performed as described in Experiments 1 and 2.

### Gain Modulation Analysis

To examine the influence of reward (Experiments 1 and 3; n = 77), errors (Experiments 1-3; n = 106), and outcome entropy (Experiments 1-3; n = 106) on the gains of the PID terms, we re-ran our PID regression analysis, including interaction terms for each type of gain modulation. In Experiment 1, the reward modulator consisted of binary reward feedback that was given on a random subset of trials, conditional on participants’ error being within a pre-specified threshold. This feedback was not correlated with absolute error on the task. In Experiment 3, the reward modulator was the number of points that participants received on each trial, which was a time-varying non-linear function of absolute error (see procedure above). In this task, participants received both error and reward feedback on every trial. Absolute error was correlated with the reward (median *r* = -.68) but Belsley collinearity diagnostics (Belsley, Kuh, & Welsch, 1980) indicated that the collinearity between absolute error and reward was below standard tolerances, suggesting that our regression would be able to assess the independent contributions of each factor. In all three experiments, the error modulator was the absolute prediction error. Outcome entropy was defined as the natural logarithm of the outcome sample standard deviation over the current and 10 previous trials within each block (with a truncated window for the first 10 trials in each block).

A robust regression (bisquare weighted) was run for every participant in every experiment, excluding the reward modulator for Experiment 2. The regression model included all main effects, as well as the interactions between the PID terms and gain modulators (*u* ~ 1 + (P + I + D)*reward + (P + I + D)*absolute error + (P + I + D)*outcome entropy). We mean-centered betas within their respective experiment and then re-centered the betas on their grand mean, removing between-experiment variance (Cousineau, 2005).

### Kalman Filter Analysis

Our Kalman filter analysis was based on the algorithm used in Kording et al (2007), building off the code that accompanied their publication. This Kalman filter model estimated the likelihood of different states using an uncertainty-weighted delta-rule algorithm. Each state was a differential equation which defined a random walk over a specific timescale (i.e., slowly-changing or quickly-changing outcome locations). See Kording et al (2007) for a complete description of this algorithm. While the Kalman filter is not optimized for our task, given that the outcomes were not generated from a random walk, it has nevertheless proved to be a good model of behavior in previous experiments that used a random walk generative function (e.g., Kakade & Dayan, 2002; Daw et al., 2006; Kording et al., 2007; Gershman, 2015).

Following Kording et al. (2007), states were defined as 30 logarithmically spaced diffusion timescales between 2 trials and the length of the experiment. We fit state noise parameters for each participant using restricted maximum likelihood estimation (MATLAB’s fmincon). The initial mean was set to the first outcome, the initial covariance was set to a small variance constant (10^−^4). As in our PID analysis, we fit the Kalman filter’s parameters so as to minimize the difference between its prediction updates and each participants’ prediction updates, based on participants’ errors on each trial (i.e., one-step look ahead).

We also compared the PI model against a variant of the Kalman filter that is less commonly used to describe adaptive behavior, but was better suited for our experiment. This Kalman filter tracks randomly drifting changes in both the position (*x*) and velocity (*ẋ*) of the outcome locations:

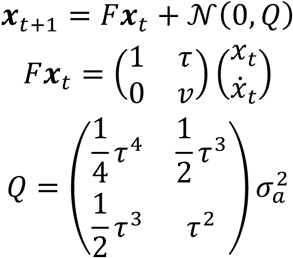

We used restricted maximum likelihood estimation to fit participant-specific velocity decay (*v*), time delay (*τ*), and state noise (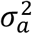) parameters to participants’ updates using the same one-step look ahead procedure described above. The initial mean was set to the first outcome, and the initial covariance was set to the variance in outcome position and velocity, averaged across participants.

## Results

### Model-Agnostic Analysis

Regressing the current and ten leading errors onto the current update (see Experiment 1 Methods), we replicated the observation that participants were influenced by past errors (Mean(SD) summed betas: 0.23(0.21), *p* ≤ 10^−5^; see Fig. 5A). Our model-generated behavior again showed that the delta-rule model categorically fails to capture the influence of leading errors. Unlike Experiment 2, here we found that the weighting of previous errors was best fit as a linear decay from the current trials, resembling PI control (Mean(SD) trend beta: linear = 0.0087(0.016), *p* < 10^−4^; quadratic = −0.002(0.016), *p* = .39; sign-randomization test). This discrepancy from Experiment 2 may be because rewards in Experiment 3 were highly dependent on accuracy. This may have biased participants more toward integral control (which favors accuracy) and away from derivative control (which favors stability; Aström & Murray, 2010).

**Figure 5.**
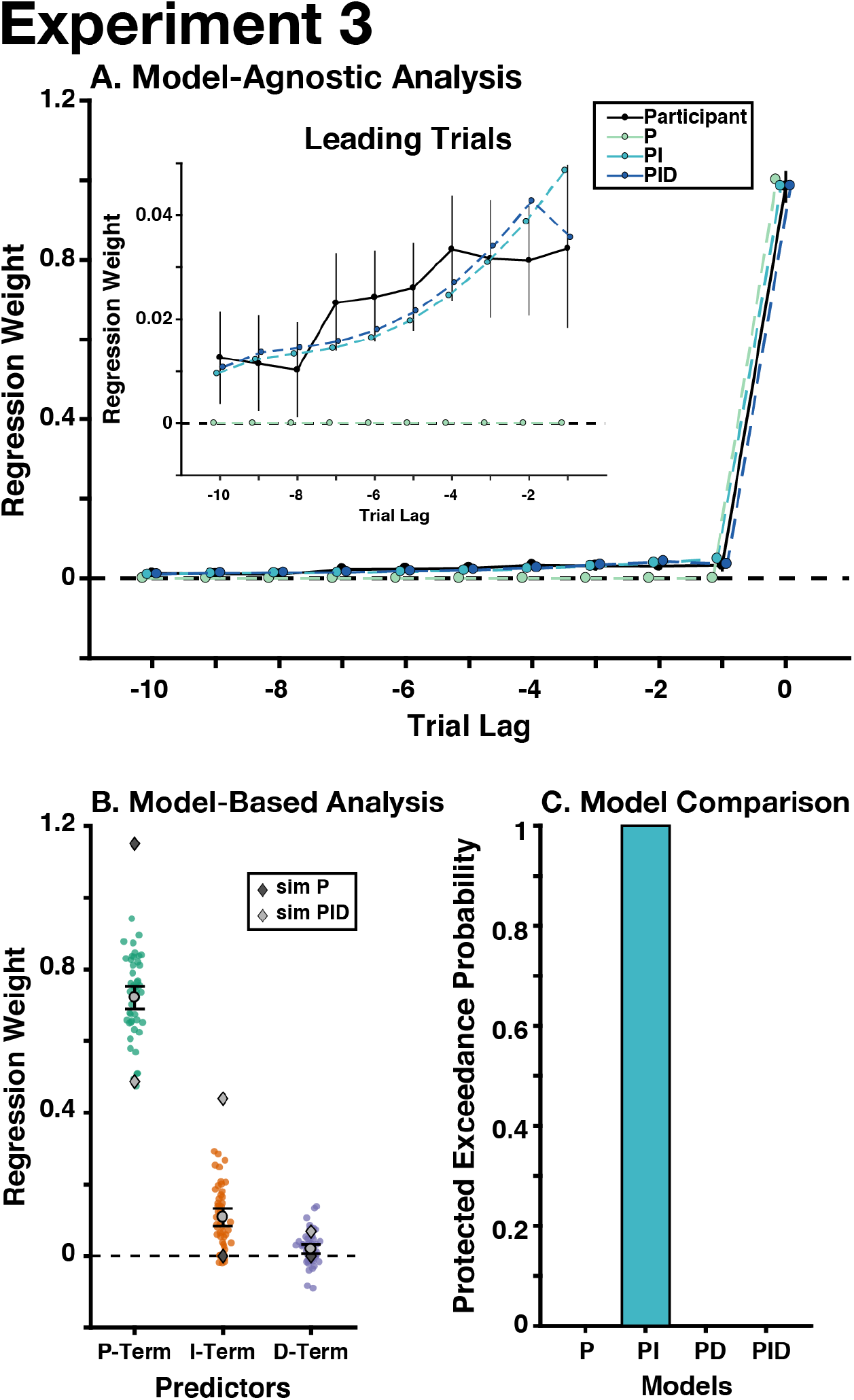
Experiment 3 Results. **A)** Participants adjusted their choices based on previous errors, unlike the predictions of a proportional controller (i.e., delta-rule model). **B)** P, I, and D control significantly predicted participants’ updates. **C)** The PI controller best explained participants’ behavior. See Figure 3 for detailed graph legends. Error bars indicate mean and between-participant bootstrapped 95% confidence intervals.

### PID Model Fit

Replicating Experiments 1 and 2, we found that our standard PID model accounted for most of the variance in participants’ updates (median *R*^2^ = 0.81). The parameters for the P-, I-, and D-terms were all significantly different from zero (Mean(SD) standardized beta; *β_P_* =0.72(0.11), *p* ≤ 10^−^5; *β_I_* = 0.11(0.086), *p* ≤ 10^−^5; *β_D_* = 0.020(0.047), *p* = .006; Fig. 5B). The group level λ was .8016. Participants’ estimated gains were similar to the ideal PID controller, but they over-weighted proportional control and under-weighted integral control. We found that there were likely differences between the model likelihoods (Bayesian Omnibus Risk < .001), and that Bayesian Model Selection strongly favored the PI model (PXP_PI_ > 0.99) over the alternate models (all other PXPs < 10^−^4; Fig. 5C).

### Gain Modulation

We examined the independent influence of rewards, absolute error, and outcome entropy in modulating the PID gains across our three experiments. We found that all three modulators significantly interacted with the P, I, and D Terms, but in distinct ways (Fig. 6): increased reward led to an increased P and I gain, and a decreased D gain (Fig. 6A; Mean(SD) interaction beta: β *P:reward* = 0.086(0.12), *p* ≤ 10-5; β*I:reward* = 0.0098(0.036), *p* = .016; β*D:reward* = −0.010(0.040), *p* = .032; sign-randomization test). Increased absolute error led to an increased P gain, and a decreased I and D gain (Fig. 6B; Mean(SD) interaction beta: β*P:error* = 0.043(0.078), *p* ≤ 10-5; β *I:error* = −0.032(0.053), *p* ≤ 10-5; β*D:error* = −0.014(0.062), *p* = .019); Increased outcome entropy led to a decreased P gain, and an increased I and D gain (Fig. 6C; Mean(SD) interaction beta: A control theoretic model of adaptive behavior 22 β*P:entropy* = −0.018(0.056), *p* = .0016; β*I:entropy* = 0.057(0.059), *p* ≤ 10-5; β*D:entropy* = 0.0098(0.039), *p* = .011).

**Figure 6.**
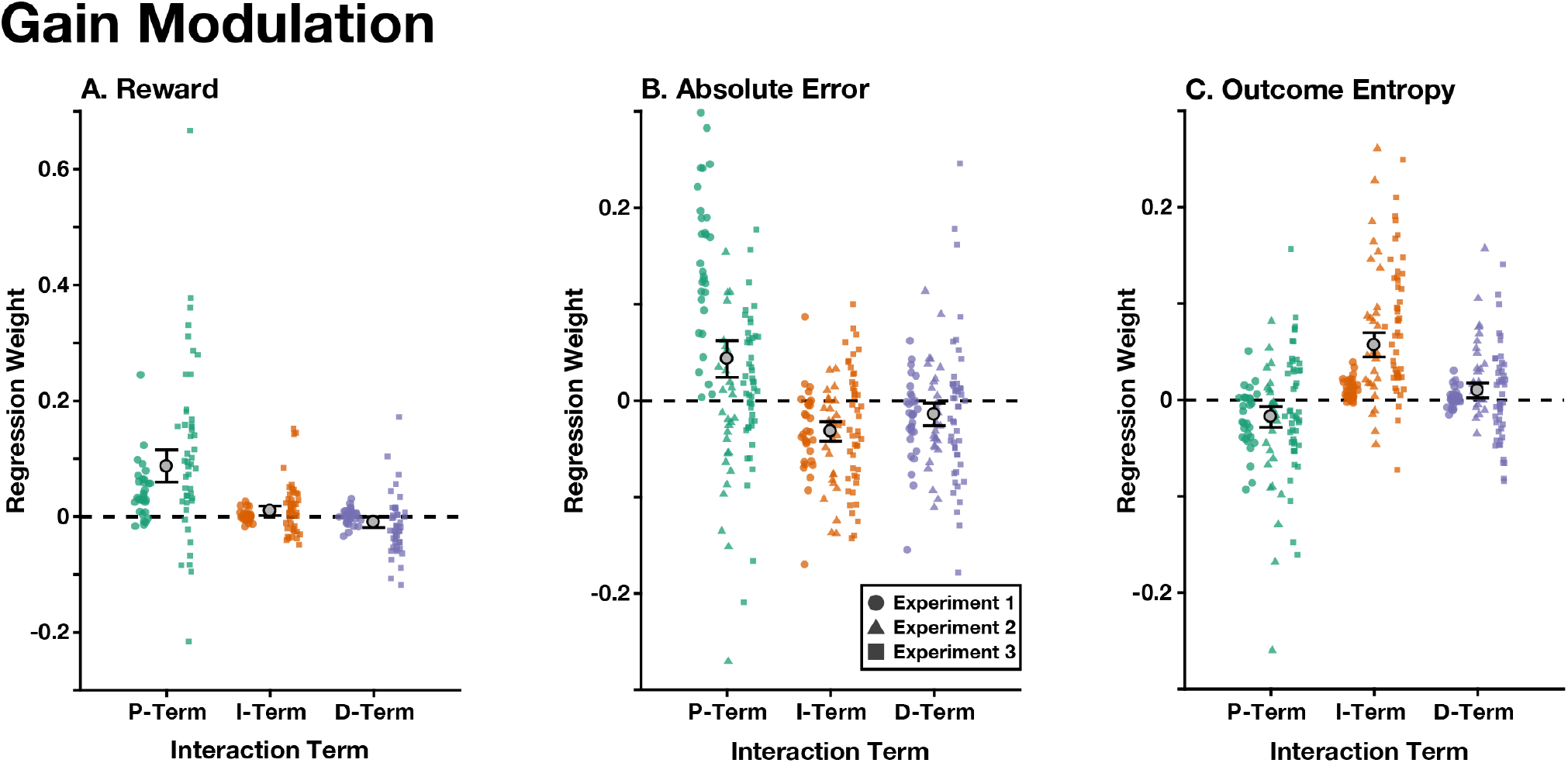
Gain Modulation. Interaction terms between the trial-wise reward (**A**), absolute error (**B**), and outcome entropy (**C**) with each of the PID terms, for each of the experiments. All models included all main effects. Colored shapes indicate individual participant’s standardized betas in each experiment (see legend). Error bars indicate mean and between-participant bootstrapped 95% confidence intervals, uncorrected for between-experiment variance.

These interactions were robust to several quality checks. First, all effects remained significant when we corrected for multiple comparisons using the Holm–Bonferroni procedure (Holm, 1979). Given the presence of outliers, we also tested our effects using a robust Wilcoxon signed-rank test (Wilcoxon, 1945), finding that all interactions remained significant (*p*s ≤ .014). Finally, we also found that all interactions remained significant when we did not remove between-experiment variance (*p*s ≤ .035; Fig. 6 depicts participants’ raw interaction betas).

### Kalman Filter Analysis

We fit the Kalman filter to participants’ behavior in both Experiments 2 and 3, finding that Bayesian model selection strongly favored the PI control model over the standard Kalman filter (pooling across experiments; PXP_PI_ > 0.99, BOR < 10^−^14; Fig. 7A). Using our lagged regression analysis approach, we also found that the standard Kalman filter’s updates depended on previous errors in a qualitatively different way from participant updates. Unlike participants the Kalman filter placed negative weights on errors made in earlier trials (Fig. 7B). We also found that the standard Kalman filter also performed especially poorly when outcomes changed over time (i.e., at different outcome velocities), whereas participants and the PI model were able to accommodate such changes in outcomes (Fig. 7C).

**Figure 7.**
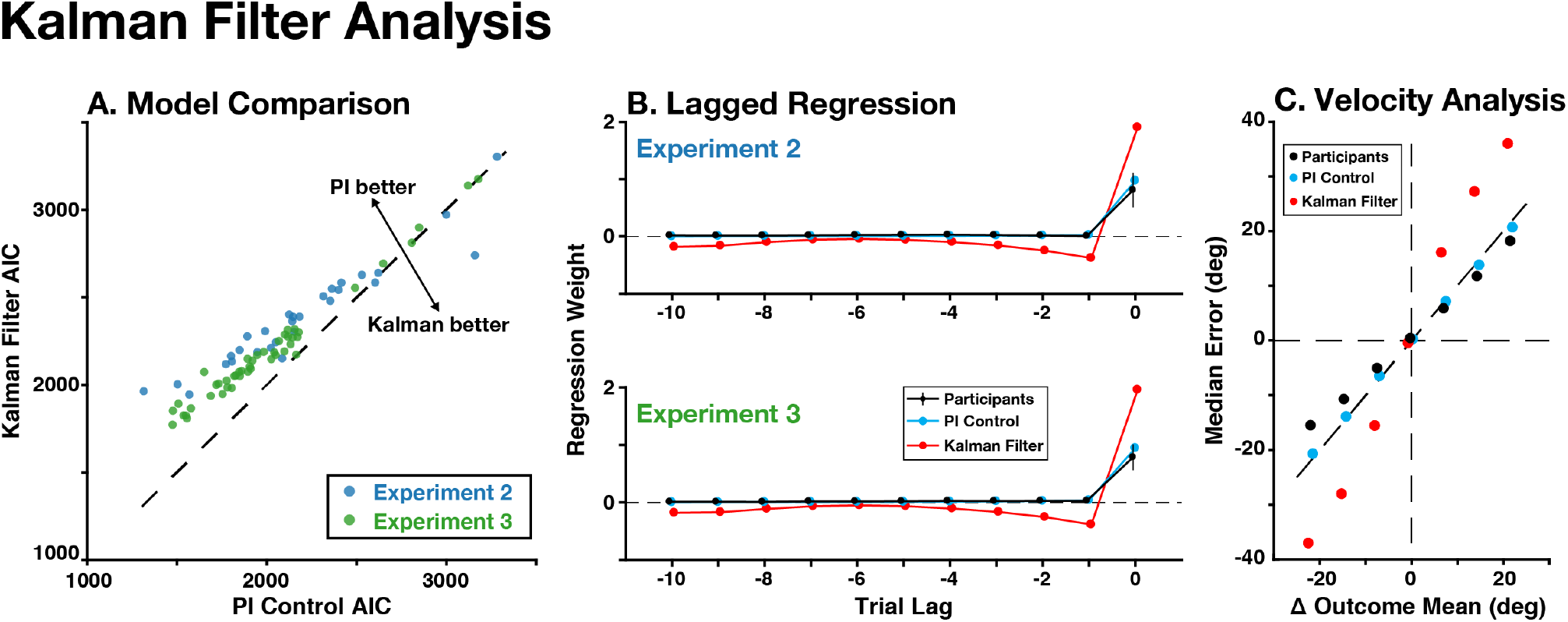
Kalman Filter Analysis. The PI control model fit participants’ behavior better than the standard Kalman filter model. **A)** For most participants (colored dots), a complexity-corrected measure of fit (AIC) was better for the PI control model than the Kalman filter model. **B)** Unlike the PI control model, the Kalman filter model poorly resembled how lagged errors influenced participants’ updates (compare to panel A in Figs 4-5). **C)** Unlike the PI control model, the Kalman filter did not resemble participants’ accuracy when the outcome distribution changed over time (pooled across Experiments 2 and 3).

We also compared the PI control model against a Kalman filter model that tracked the position and velocity of outcomes over time. Despite the additional complexity of this model, we found that the PI model fit similarly well (PXP_PI_ = .63, BOR = .70). These models were identifiable, as we could accurately recover the correct model when either of them generated behavior, suggesting that they offer dissociable explanations of participants’ behavior. Interestingly, we found that participants’ velocity estimates strongly decayed over time (mean *v* = 0.32), and that this parameter strongly correlated with participants’ integral gain (*r* = .79, *p* < 10^−^16), suggesting that these terms might serve complementary computational roles. Collectively, these results show that the PI model offers a more parsimonious account of participants’ behavior than complex, task-informed inferential models.

## Discussion

In Experiment 3, we found confirmatory evidence that the PI model accurately describes participants’ predictions, and that participants adjust their weighting of different PID terms based on trial-wise task dynamics. We found that each PID term was uniquely sensitive to changes in reward, absolute error, and outcome entropy, extending previous observations of the role of these modulators on proportional control, and providing further evidence that the PID terms represent distinct control processes. We also found that the PI model offered a better explanation of behavior than the standard Kalman filter, and performed similarly to a specialized Kalman filter variant, demonstrating that the PI model is as powerful as more complex models based on explicit state space representations.

When participants received larger rewards, they modulated their gains in a way that is consistent with a preference for accuracy (P- and I-Terms) over stability (D-term; Ang, Chong, & Li, 2005), and potentially indicating an exploitive strategy for the high reward environments (Kovach et al., 2012). While participants’ proportional gain was already larger than the best-performing gain, this may reflect the unique role of reward modulation, when controlling for the environmental changes (e.g., entropy) that make a high proportional gain less desirable. Another alternative is that the P- and/or I-Terms are effortful to implement, with rewards ‘paying the cost’ of these control policies (Kool, Gershman, & Cushman, 2017; Manohar et al., 2015). Further work will be necessary to dissociate the role of salience and motivation on reward-modulated gain adjustments.

In response to absolute errors (i.e., surprise), participants increased their immediate adjustment (P-Term), and relied less on previous feedback (I- and D-Terms). This is consistent with the idea that large errors may indicate changes in the environment (Pearce & Hall, 1980; Nassar et al., 2010), and with filtering mechanisms in industrial PID control that improve robustness by limiting the long-term influence of noisy samples (Ang et al., 2005).

While outcome entropy, and uncertainty, has traditionally been thought to increase the gain on proportional control (Behrens et al., 2007; Courville et al., 2006; Nassar et al., 2010), in our experiment the P-Term was decreased, and the I- and D-Terms were instead increased. Interestingly, when we instead implement this gain modulation in a P-only model, we do find that outcome entropy increases the gain of the P-term (data not shown). Unlike previous experiments studying uncertainty, environmental change in Experiments 2 and 3 required tracking gradually changing outcomes, which accounted for most of the outcome entropy and for which integral and derivative control are particularly useful (Kovach et al., 2012; Wittmann et al., 2016). In Experiment 1, where these gradual changes were not present, we found that uncertainty after a change-point increased control gains (McGuire et al., 2014), which may be reflected here by integrating over the trials since the change-point.

We found that the PI model explained participants behavior better than the standard Kalman filter (a powerful model of adaptive learning; Kording et al., 2007), and that the Kalman filter failed to capture participants’ use of feedback history. This difference was largely due to the ability of the integral term to track constant drifts in the environment, epochs that were poorly accounted for by the Kalman filter. Interestingly, the Kalman filter’s updates were negatively correlated with errors made on earlier trials (when controlling for the influence of the current error). We believe that this is due to the fast diffusion timescales, which were updated the fastest (they were set to the highest state noise, as in Kording et al., 2007), and define the difference between current and recent trials. We found that the lagged influence of recent trials was more strongly negative for shorter timescales (data not shown).

We also compared the PI model against a specialized Kalman filter that tracked both the position and velocity of outcomes, finding that these models fit similarly well. There was a strong relationship between the Kalman filter’s velocity term and the PI controller’s integral term, suggesting that participants could use integral control to track constant drifts in the environment. This position-velocity Kalman filter has received little attention in the learning literature and warrants further investigation, however, it currently offers a less parsimonious explanation of behavior than PI control due to its greater computational complexity and its requirement for explicit state representations. While these Kalman filters did not offer better models than PI control, the Kalman filter embodies the same principles as our adaptive gain analysis: control gains should be adaptive and depend on factors like environmental stability.

## General Discussion

Across three experiments, we found that the PI model successfully captured participants’ prediction updating in a stochastic environment. By incorporating a richer model of control monitoring and adjustment, the PI controller was able to account for ways in which performance in such environments deviates from predictions of standard error-driven (delta-rule) learning models. We also replicated and extended previous findings showing that learning parameters themselves are modulated by environmental signals (e.g., reward), and extended these findings to show that signals related to the magnitude of reward, error, and outcome entropy can differentially affect the gains on the PID model parameters.

Our findings suggest that PI control offers a good account of behavior across two fairly different task environments. Indeed, while we found that normative PID gains differed substantially between Experiment 1 (discrete transitions) and Experiments 2-3 (gradual transitions), participants’ behavior continued to qualitatively match the behavior predicted by this normative controller across studies, in each case matching the sign of the best-performing control gain. This suggests that these gains adapted to the specific environment that participants were acting in. Specifically, when outcomes were prone to shift sharply and dramatically (Experiment 1), participants tended to rely less on history-dependent control processes like integral and derivative control, especially on trials in which large errors may have indicated a state shift.

While we have focused our discussion of the PID controller on all three of its components, in industrial settings the D-Term is often given the lowest gain or not included (Aström & Murray, 2010), as it is highly sensitive to noise. Accordingly, our own data supported little to no role for the derivative term in the current experiments, both normatively and in our model fits to participants’ behavior. While the derivative control term was significant in all of the experiments, and interacted with the absolute error, it did not account for sufficient variance to outweigh complexity penalties in model comparison. This may have been compounded by the fact that the derivative term was negatively modulated by absolute error, which may have caused it to explain less of the variance on trials where there were large updates. While the outcomes in Experiments 2 and 3 were designed to differentiate PID control from the delta-rule model, they were not designed to specifically detect derivative control. Future research should investigate cases where derivative control is especially beneficial for good performance. Since derivative control provides high-frequency compensation to improve responsivity, it may be the case that derivative control is generally poorly suited for tasks that depend on inter-trial adjustments and favor accuracy over speed. Relative to Experiment 2, Experiment 3 emphasized accuracy through its reward structure, and deemphasized responsivity due to its longer trial length. While there were several differences between these experiments, these factors may have contributed to the differences in derivative control between these experiments.

Some of the most promising evidence for derivative control was that in Experiment 2 participants down-weighted recent errors (from *t*-3 and *t*-1) relative to what would be expected by error integration alone. While basic derivative control would only compare the current and previous errors, participant’s behavior resembles a common practice in control engineering to low-pass filter the derivative term to improve robustness (Ang et al., 2005). The discrepancy between the observed non-linear influence of previous errors (predicted by the full PID model) and the model selection preference for the PI model, may therefore be accounted for by alternative forms of derivative control.

We found that the PID terms were independently modulated by reward (Experiments 1 and 3), absolute error (Experiments 1-3), and outcome entropy (Experiments 1-3) on a trial-to-trial basis. While there is substantial literature on how environmental factors should influence the standard delta-rule model, less is known on how these factors should modulate PID gains. While we have proposed speculative explanations for the role of each modulating factor, at a minimum, the unique pattern of interactions for each of the PID terms suggests that P, I, and D represent dissociable forms of control. Future experiment should examine the extent to which gain modulation depends on the structure of the task and environment, for instance whether the task rewards consistency in addition to accuracy.

The PID model provides robust control without relying on an explicit model of the environment, offering a parsimonious explanation of the behavior we observed in these experiments. However, there have been notable successes for algorithms that instead learn generative models of the environment (e.g., using Bayesian updating), and can represent the uncertainty about upcoming choices (e.g., Daw, Niv, & Dayan, 2005; Franklin & Frank, 2015; Griffiths, Lieder, & Goodman, 2015; McGuire et al., 2014; Nassar et al., 2010, though see Geana & Niv, 2014; Duverne & Koechlin, 2017). To examine this possibility, we compared the PI control model against the Kalman filter, a standard model for state estimation in the face of noise and uncertainty. We found that the PI model better explained participants’ behavior than the standard Kalman filter, and fit comparably to a Kalman filter that was specialized for this experiment. In contrast to the Kalman filter, the PID controller offers a general control process that can parsimoniously account for participants behavior with minimal knowledge about the task structure. These benefits would likely be compounded by the complex dynamics of natural environments.

Despite these promising results, we would not rule out the possibility that participants rely on a combination of both model-free (e.g., PID) and model-based control (Gläscher, Daw, Dayan, & O’Doherty, 2010; Daw, Gershman, Seymour, Dayan, & Dolan, 2011; Kool, Cushman, & Gershman, 2007; Momennejad et al., 2017; Korn & Bach, 2018). Previous experiments have demonstrated the utility of model-based predictions for explaining participants’ behavior in other environments, and participants can report confidence in their choices. Model-based control may serve to modulate the PID controller itself (e.g., to tune gain parameters or reset control processes; Bouret & Sara, 2005; Behrens et al., 2007; Nassar et al., 2010; McGuire et al., 2014); may be selectively engaged in environments that are stable, constrained, or familiar; and/or may trade-off over different stages in learning (Denève, Alemi, & Bourdoukan, 2017).

Another promising feature of the PID model is that it offers a model of behavioral control that can be plausibly implemented by a neural system. There have been several neural network implementations of PID controllers in industrial engineering (e.g., Cong & Liang, 2009), with integral and derivative control implemented as positive and negative recurrent circuits, respectively. This simple architecture demonstrates the ease with which a neural system could develop PID control dynamics. Moreover, recent studies have found neuroscientific evidence that is broadly consistent with the predictions of such an architecture. For instance, Bernacchia and colleagues (2009) found that in rhesus macaques’ cingulate and prefrontal cortices, large populations of neurons encoded the history of trial-epoch-selective activity, likely including error-related responses (cf. Seo & Lee, 2007). Each of these regions contained equally-sized populations of neurons that tracked either the exponentially-weighted sum of recent trials, or the difference between recent and previous trials, putative markers of integral and derivative control, respectively. Convergent data in humans found that fMRI activity in dorsal ACC parametrically tracked a recent history of prediction errors in a changing environment (Wittmann et al., 2016), again consistent with the operations of an integral-based controller. Accordingly, these authors found that incorporating integration into their behavioral model explained choices in their task better than the traditional delta-rule model. While these findings provide evidence for neural signatures of feedback history (see also: Kennerley et al., 2006; Seo & Lee, 2007), and are consistent with the monitoring function of PID control, future experiments are needed to formally test for the neural correlates of this model.

These experiments together provide strong evidence for the viability of control theoretic models as mechanisms of human prediction updating in dynamic environments. This class of models has been highly influential in research on motor control, including the PID controller in particular (e.g., Kawato & Wolpert, 1998). Motor control models typically describe the rapid regulation of limb movements to produce trajectories that are fast, accurate, and robust. In contrast, participants in our experiments were not motivated to make fast or accurate trajectories, and instead may have used analogous control process to adapt their predictions from trial to trial. Control theoretic algorithms (like PID control) may be a domain-general class of neural functions, involved in a diverse array of cognitive processes (Ashby, 1956; Powers, 1973; Pezzulo & Cisek, 2016), including the cognitive control functions that have been suggested to operate using both classical (Botvinick et al., 2001) and optimal (Shenhav et al., 2013) control principles. The architecture of these executive control algorithms, and the nature of the references that they regulate, are important areas of further research.

## Acknowledgments

We would like to thank Kia Sadahiro and William McNelis for their assistance in data collection.

## Data Availability

All data and analysis scripts are available upon request.

## Footnotes

1 Domain-specific ‘delta-rule’ algorithms are common in many fields, such as the *Rescorla-Wagner learning rule* (Rescorla & Wagner, 1972), or a delta rule algorithm used in neural networks (Widrow & Hoff, 1960). In this paper, we define the delta-rule as a more general class of error-based learning rules in which adjustments are proportional to errors.

2 We chose AIC over the more conservative Bayesian Information Criteria (BIC) because model recovery found that BIC was overly conservative: model selection using BIC did not prefer the full PID model when this model generated behavior (i.e., when PID was the ground truth). While AIC is not the ideal fit metric for Bayesian model selection (as it is not an approximation of model likelihood), the development team for SPM’s Bayesian model selection protocol have justified using AIC as a legitimate alternative to BIC: ‘*Though not originally motivated from a Bayesian perspective, model comparisons based on AIC are asymptotically equivalent to those based on Bayes factors (Akaike, 1973a), that is, AIC approximates the model evidence.’* (Penny et al., 2004, *p*. 1162; see also: Penny, 2012; Rigoux et al., 2014).

